# A Two-Part Strategy using Genomic Selection in Hybrid Crop Breeding Programs

**DOI:** 10.1101/2020.05.24.113258

**Authors:** Owen Powell, R. Chris Gaynor, Gregor Gorjanc, Christian R. Werner, John M. Hickey

## Abstract

Hybrid crop breeding programs using a two-part strategy produced the most genetic gain, but a maximum avoidance of inbreeding crossing scheme was required to increase long-term genetic gain. The two-part strategy uses outbred parents to complete multiple generations per year to reduce the generation interval of hybrid crop breeding programs. The maximum avoidance of inbreeding crossing scheme manages genetic variance by maintaining uniform contributions and inbreeding coefficients across all crosses. This study performed stochastic simulations to quantify the potential of a two-part strategy in combination with two crossing schemes to increase the rate of genetic gain in hybrid crop breeding programs. The two crossing schemes were: (i) a circular crossing scheme, and (ii) a maximum avoidance of inbreeding crossing scheme. The results from this study show that the implementation of genomic selection increased the rate of genetic gain, and that the two-part hybrid crop breeding program generated the highest genetic gain. This study also shows that the maximum avoidance of inbreeding crossing scheme increased long-term genetic gain in two-part hybrid crop breeding programs completing multiple selection cycles per year, as a result of maintaining higher levels of genetic variance over time. The flexibility of the two-part strategy offers further opportunities to integrate new technologies to further increase genetic gain in hybrid crop breeding programs, such as the use of outbred training populations. However, the practical implementation of the two-part strategy will require the development of bespoke transition strategies to fundamentally change the data, logistics, and infrastructure that underpin hybrid crop breeding programs.

**Key message:** Hybrid crop breeding programs using a two-part strategy produced the most genetic gain by using outbred parents to complete multiple generations per year. However, a maximum avoidance of inbreeding crossing scheme was required to manage genetic variance and increase long-term genetic gain.

## Introduction

The two-part strategy produced the most genetic gain in hybrid crop breeding programs, but the maximum avoidance of inbreeding crossing scheme was required for it to increase long-term genetic gain. The two-part strategy uses outbred parents to complete multiple generations per year in hybrid crop breeding programs. In contrast, conventional plus genomic selection strategies are limited in this regard by the time they take to develop inbred lines. The maximum avoidance of inbreeding crossing scheme manages genetic variance by maintaining uniform contributions and inbreeding coefficients across all crosses. This study performed stochastic simulations to quantify the potential of a two-part strategy to increase the rate of genetic gain in hybrid crop breeding programs. A large increase in food production is required to meet the demand for a global population of 9 billion people in 2050. Increasing the rate of genetic gain of breeding programs is one route to achieve sustainable, permanent and cumulative increases in food production. Hybrid crops, and their genetic improvement, have made major contributions to historical increases in food production. For example, genetic merit for maize yield has approximately doubled from 1930 to 2001 (Duvick et al. 2010; Fig. 4.1). However, the current rates of genetic gain in hybrid crop breeding programs are insufficient to meet the estimated 70% increase in overall food production (Alexandratos and Bruinsma 2012) required within the next 30 years.

Genomic selection could increase the rate of genetic gain in hybrid crop breeding programs by directly addressing three of the parameters of the breeder’s equation. The breeder’s equation provides a framework to understand how the rates of genetic gain in breeding programs can be increased (Lush 1943). The breeder’s equation shows that genetic gain is a function of (i) the accuracy of ranking selection candidates based on genetic merit, (ii) the intensity of selection, (iii) the genetic variance in the population, and (iv) the generation interval. Genomic selection could increase the rate of genetic gain in hybrid crop breeding programs by reducing the generation interval, increasing the selection intensity and increasing the selection accuracy.

Hybrid crop breeding program designs involve multiple stages with selection candidates evaluated against an increasing number of testers in an increasing number of environments. As the accuracy of evaluation increases through the stages of evaluation, individuals are recycled by reciprocal recurrent testcross selection (Bernardo, 2014; Hull, 1945) and crossed to create the next set of selection candidates. Reciprocal recurrent testcross selection aims to improve the general combining ability of selection candidates. However, the development of inbred individuals in reciprocal recurrent testcross selection requires time which results in longer breeding cycle times and slows the rate of population improvement. For example, hybrid crop breeding program designs typically have a cycle time of 3 to 4 years and are not radically different from a typical breeding program design for inbred crops.

Genomic selection can increase the rate of population improvement in plant breeding programs. Recently, Gaynor et al. (2017) proposed a two-part breeding strategy for inbred crops that explicitly separates a conventional plant breeding program into two distinct components. These components are:

i. a population improvement component to develop improved germplasm via recurrent genomic selection, and;
ii. a product development component to identify new inbred varieties within conventional plant breeding program designs.

Gaynor et al. (2017) used simulation to compare conventional and two-part breeding program designs in the context of inbred crops. Compared to the conventional design, the two-part strategy generated 2.4 times more genetic gain per unit cost and unit time.

Conceptually the two-part strategy is equally suited to hybrid crop breeding programs. In a hybrid crop breeding program, the two-part strategy could enable large increases in genetic gain by shortening the generation interval considerably. However, in the context of hybrid crops, population improvement would need to be driven by reciprocal recurrent genomic selection rather than recurrent genomic selection. Reciprocal recurrent selection aims to improve the general combining ability of individuals in different heterotic pools. Reciprocal recurrent genomic selection (Kinghorn et al. 2010) uses the phenotypes and parental genotypes of hybrids to more accurately estimate the general combining ability of individuals (Rembe et al. 2019).

However, the previous implementations of the two-part strategy in inbred crops showed that large reductions of the generation interval came at the expense of genetic variation. Gaynor et al. (2017) showed that two-part breeding programs that used rapid cycling to reduce the generation interval below 0.5 years reduced long-term genetic gain. Using simulation, Gorjanc et al. (2018) showed that long-term genetic gain can be optimised with crossing schemes that balance increases in genetic gain with reductions in genetic variance. Maximum avoidance of inbreeding is a crossing scheme that maintains uniform contributions across generations and uniform inbreeding coefficients across all crosses (Wright 1921; Kimura and Crow 1963). Due to the large number of generations per year that can be completed with the two-part strategy, the use of maximum avoidance of inbreeding could increase long-term genetic gain in hybrid crop breeding programs.

The objective of this study was to develop and test the two-part strategy in the context of hybrid crops. Stochastic simulations were used, with a maize breeding program as a model, to compare conventional, conventional plus genomic selection and two-part hybrid crop breeding programs under an assumption of approximately equal operating costs and time. To manage long-term genetic variance, both a circular crossing scheme and a maximum avoidance of inbreeding crossing scheme were used in conjunction with genomic selection. The results show that: (i) the implementation of genomic selection in hybrid crop breeding programs increases the rate of genetic gain, (ii) the two-part strategy was the most cost-effective strategy for implementing genomic selection in hybrid crop breeding programs, and (iii) two-part hybrid crop breeding programs completing multiple selection cycles per year should use methods to manage genetic variance.

## Methods

Stochastic simulations of entire hybrid crop breeding programs were used to compare:

- a conventional breeding program not using genomic selection;
- three conventional plus genomic selection breeding programs, and;
- two breeding programs implementing the two-part strategy.

These breeding programs were compared on an equal time across 40 years of breeding. Each breeding program was constrained to have approximately equal operating costs so that direct comparisons between the different breeding programs would represent their relative effectiveness. The six different breeding programs were compared using 10 independent replicates of a stochastic simulation for three levels of genotype-by-year interaction variance. Each replicate consisted of:

i. a burn-in phase shared by all strategies so that each strategy had an identical, realistic starting point, and;
ii. a future breeding phase that simulated 20 years of future breeding with each of the different breeding strategies.

### Burn-In Phase

Specifically, the burn-in phase was subdivided into three stages. The first stage simulated the species’ genome sequence. The second stage simulated trait architecture and founder genotypes for the initial parents. The third stage simulated 20 years of breeding using the conventional breeding strategy without genomic selection.

#### Generation of whole genome sequence data

For each replicate, a genome consisting of 10 chromosome pairs was simulated to resemble the maize genome. These chromosomes were assigned a genetic length of 2.0 Morgans and a physical length of 2×10^8^ base pairs. Sequences for each chromosome were generated using the Markovian Coalescent Simulator (Chen et al. 2009) within AlphaSimR (Gaynor et al. 2019). Recombination and mutation rates were respectively set to 1.25×10^−8^ per base pair and 1×10^−8^ per base pair. Historical effective population size was simulated, beginning with a single population, as follows; 100,000 at 6,000 generations ago, 10,000 at 2,000 generations ago, 5,000 at 1,000 generations ago, 1,000 at 100 generations ago. To mimic the genetic separation of two heterotic groups the population was split 200 generations ago. The final effective population size at the end of the coalescent simulation was set to 100 for each heterotic group. These values were chosen to roughly follow the evolution of effective population size in North American hybrid maize.

#### Founder Genotypes

The founders served as the initial parents in the burn-in breeding phase. This was accomplished by randomly sampling gametes from the simulated genome to assign as sequences for the founders. 80 founders were created for each of the two heterotic groups. Sites segregating in the founders’ sequences were randomly selected to serve as 2000 single nucleotide polymorphism (SNP) markers per chromosome (20000 total) and 300 quantitative trait nucleotides (QTL) per chromosome (3000 total). The randomly selected sites for SNP markers and QTL were not allowed to overlap. The founders were converted to inbred lines by simulating the formation of doubled haploids (DH).

#### Trait Architecture

Three types of biological effects were modelled at each QTL to simulate genetic values: additive effects, dominance effects and genotype-by-year effects. Under the AlphaSimR framework, this is referred to as an ADG trait. We will give only a brief summary of the modelling procedure, while a detailed description can be found in the vignette of the AlphaSimR package (Gaynor et al. 2019).

A single trait representing grain yield, controlled by 3,000 QTL, was simulated for all individuals. Each QTL was assigned an additive genetic effect, composed of additive and genotype-by-year effects, and a dominance effect resulting from the interaction between alleles at a heterozygous locus. Epistatic gene action was not considered. Three levels of genotype-by-year variance were examined: 0, 2 and 4 times the genetic variance. Dominance effects were calculated by multiplying the absolute value of each QTL allele effect by a locus-specific dominance deviation (**δ**). Dominance deviations were sampled from a normal distribution with mean dominance deviation of 0.92 and variance of 0.2, to approximate historical levels of heterosis displayed in commercial maize (Troyer and Wellin 2009). The genetic value of each QTL was then defined as the sum of additive QTL effects and the dominance effect of interacting alleles. Finally, the genetic value of each individual was obtained as the sum of all of the QTL genetic values, accounting for the individuals’ genotype at these QTL.

The genetic value of each individual was used to produce phenotypic values by adding random error. The random error was sampled from a normal distribution with mean zero. The variance of the random error varied according to the stage of evaluation in the breeding program. This was done to account for increasing accuracy in the evaluation as the number of replications per entry increased. The values for these error variances were set to achieve target levels of heritability. The levels of heritability represented heritability on an entry-mean basis for the 80 founder genotypes when genotype-by-year variance was absent. The levels of heritability that were used are presented in the description of the conventional breeding program below.

#### Conventional Breeding Program

Burn-in breeding for yield was simulated using 20 years of breeding in a conventional program without genomic selection. The design of the burn-in program approximated existing maize breeding programs taken from Bernardo (2014; Table 8.3). The key features of the two heterotic groups in this breeding program were:

i. a crossing block consisting of 80 DH lines used to develop 80 biparental populations each year;
ii. the development of 25 new DH lines from each biparental cross;
iii. a 3-year cycle time from crossing to selection of new parents; and
iv. a 6-year production interval from crossing to release of a new commercial hybrid.

All selection in the burn-in program was performed using phenotypes. These phenotypes represented direct selection on yield using a yield trial. The levels of heritability at a particular selection stage were adapted from the number of DH lines and locations reported in Bernardo (2014; Table 8.3). A schematic for the overall design of the burn-in program is given in Fig. 1 and a detailed description follows below. Each of the stages, described below, were conducted independently in the two heterotic groups. The progression of germplasm through the breeding program was simulated using AlphaSimR (Gaynor et al. 2019).

**Fig. 1.**
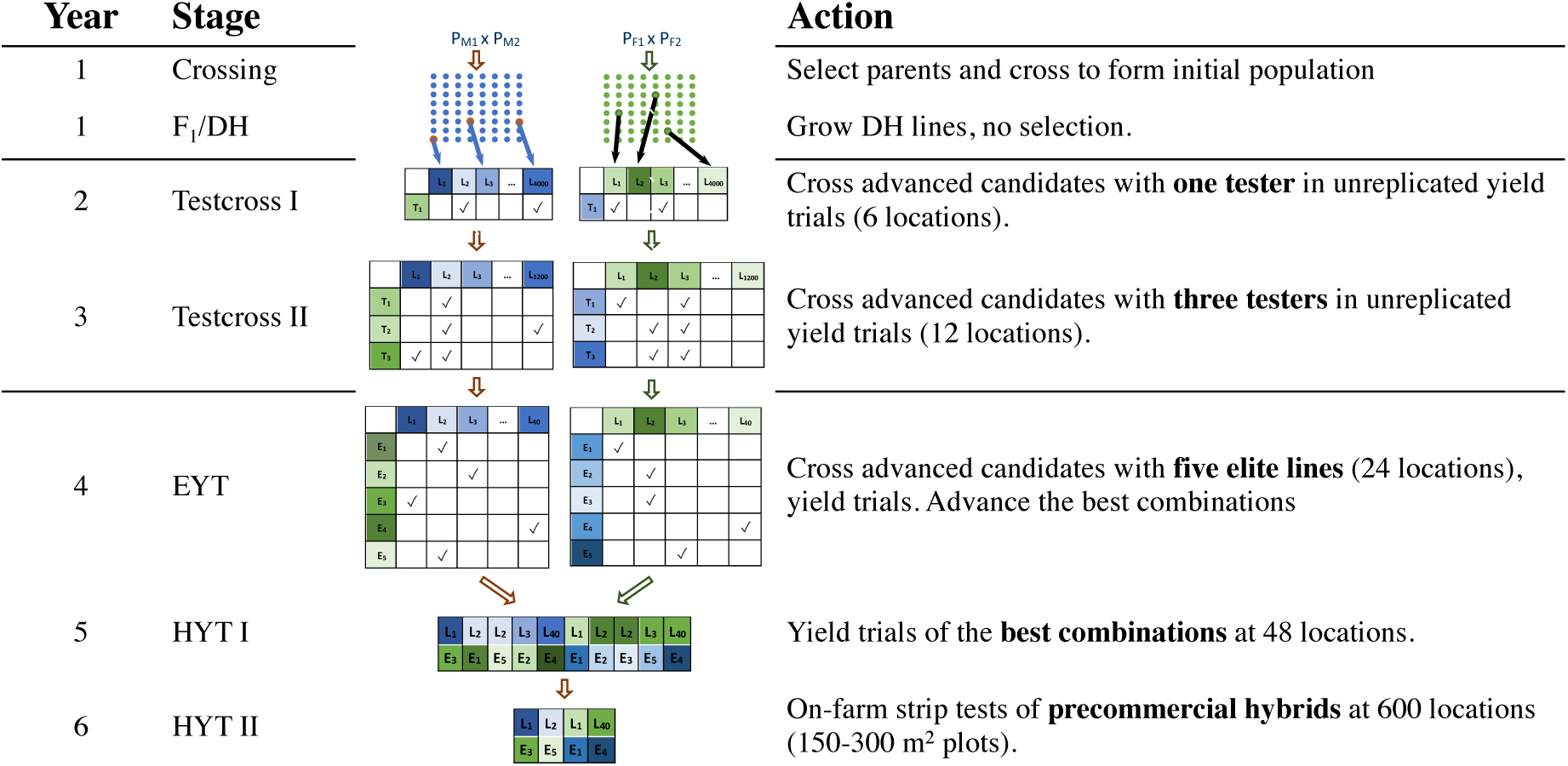
Overview of breeding schemes for the conventional hybrid crop breeding program (used in burn-in breeding) and the breeding programs using standard genomic selection strategies. DH, doubled haploid; EYT, elite yield trial; HYT I, hybrid yield trial 1; HYT II, hybrid yield trial 2

#### Year 1

80 bi-parental populations were created with intra-heterotic group crosses. It was ensured that each of the 80 parental DH lines was used as a male or female only once. Each cross produced 25 F1 derived doubled haploid lines (Geiger and Gordillo 2009). The 2,000 DH lines were planted in separate plots. No selection was performed at this stage. Each DH line was crossed to a single inbred tester.

#### Year 2

The 2,000 DH testcrosses (1 tester x 2,000 DH lines) were evaluated in the testcross 1 (TC1) stage. The TC1 stage represented yield measured in unreplicated, two-row plots across 6 locations. Selection in the TC1 stage was modelled as the selection on a yield phenotype with a heritability of 0.54. The best performing 400 DH lines were advanced to the next trial based on general combining ability. Each of the 400 DH lines was crossed to 3 inbred testers.

#### Year 3

The 1,200 DH testcrosses (3 testers x 400 DH lines) were evaluated in the testcross 2 (TC2) stage. The TC2 stage represented yield measured in unreplicated, two-row plots across 12 locations. Selection in the TC2 stage was modelled as the selection on a yield phenotype with a heritability of 0.71. The best performing 40 DH lines were advanced to the next trial based on general combining ability. These 40 DH lines were then crossed to 5 ‘elite’ DH lines from the other heterotic group. This produced 200 single cross experimental hybrids.

#### Year 4

The 200 experimental hybrids were evaluated in the elite yield trial (EYT) stage. The EYT stage represented yield measured in unreplicated, two-row plots across 24 locations. Selection in the EYT stage was modelled as the selection on a yield phenotype with a heritability of 0.82. The best performing 20 experimental hybrids were advanced to the next trial.

#### Year 5

The 20 experimental hybrids were evaluated in the hybrid yield trial 1 (HYT1) stage. The HYT1 stage represented yield measured in unreplicated, two-row plots across 48 locations. Selection in the HYT1 stage was modelled as the selection on a yield phenotype with a heritability of 0.98. The best performing 4 experimental hybrids were advanced to the next trial.

#### Year 6

The 4 pre-commercial hybrids were evaluated in the hybrid yield trial 2 (HYT2) stage. The HYT2 stage represented yield measured in on-farm strip tests of pre-commercial hybrids across 600 locations. Selection in the HYT2 stage was modelled as the selection on a yield phenotype with a heritability of 0.99. Commercial hybrids were selected for release from this set of pre-commercial hybrids.

### Future Breeding

The future breeding phase of the simulation modelled alternative breeding programs and compared them against the conventional breeding program. Each breeding program was simulated for an additional 20 years following a common burn-in breeding phase so that each strategy could be evaluated with an equivalent starting point. Two crossing schemes were used in breeding programs with a generation interval of 1 year or less. All breeding programs were constrained to equivalent operating costs (Table 1).

**Table 1.** Summary of hybrid crop breeding program sizes (number of individuals across both heterotic pools) and costs. DH, doubled haploid; EYT, elite yield trial; HYT I, hybrid yield trial 1; HYT II, hybrid yield trial 2

| **<u>Breeding Program</u>** | **<u>Parents</u>** | **<u>DH</u>** | **<u>TC1</u>** | **<u>TC2</u>** | **<u>EYT</u>** | **<u>HYT1</u>** | **<u>HYT2</u>** | **<u>Cost (\$)</u>** |
| --- | --- | --- | --- | --- | --- | --- | --- | --- |
| Conventional | 80 | 4000 | 4000 | 2400 | 400 | 40 | 8 | 1,276,800 |
| Conventional Plus Genomic Selection Strategies | 80 | 3840 | 3840 | 2196 | 400 | 40 | 8 | 1,273,680 |
| Two-Part Strategies | 80 | 3200 | 3200 | 1728 | 400 | 40 | 8 | 1,271,040 |

### Cost Equalizing Strategy

To ensure approximately equal operating costs across different breeding programs, the number of DH lines tested across the product development component was reduced in breeding programs using genomic selection. Table 1 details the exact numbers in each of the breeding programs using genomic selection. Since the two-part breeding programs used multiple cycles per year, they resulted in up to three times as many crosses per year. Therefore, the number of candidate DH lines was reduced further compared to the conventional plus genomic selection breeding programs. To equalize genotyping costs within the population improvement component of the two-part strategy, the number of genotyped offspring per cross was reduced with each additional crossing cycle per year: (i) 1 cycle/year – 80 seeds genotyped per cross; and (ii) 3 cycles/year – 26 seeds genotyped per cross. The remaining components of the breeding programs were kept constant.

Estimated costs for genotyping, phenotyping and producing DH lines were used in the equalisation of operating costs. The cost for producing doubled haploid lines was estimated at $45 based on the lowest publicly advertised price (http://www.plantbreeding.iastate.edu/DHF/DHF.htm). Genotype costs and phenotypic evaluation of a yield trial plot, were assumed to be equivalent at a cost of $15 (http://techservicespro.com/test-locations/). The cost for crossing, which in any case would be small, was not considered even though it varied between breeding programs.

### Genomic Selection Training Population & Method

Genomic selection in each hybrid crop breeding program used initial training populations comprising the last 3 years of testcross I and II yield trial data from the recent breeding burn-in phase. Separate training populations were developed for each of the two heterotic pools. Thus, the initial training populations comprised phenotypic records on 7,200 testcross genotypes. The training populations were updated in subsequent years via a 3-year sliding window approach, in which the oldest year of data was replaced with the data from the newest year. As a consequence of the cost equalisation process the training population sizes were reduced to phenotypic records on 6,858 and 5,664 testcross genotypes for conventional genomic selection and two-part breeding strategies, respectively.

The genotypes and testcross phenotypic means of the DH selection candidates were fitted using the genomic selection model used by Bernardo and Yu (2007), and the genetic background of the tester was accounted for by fitting a tester-by-stage fixed effect. A separate genomic selection model was fitted for each heterotic pool. Genomic predictions were calculated using the AlphaSimR function “RRBLUP”. This function fits a ridge regression best linear unbiased prediction model (Whittaker et al. 2000). It models the heterogeneous error variance due to different levels of error in each yield trial by weighting for the effective number of field measurements.

### Conventional plus Genomic Selection Breeding Programs

Three conventional plus genomic selection breeding programs were used to quantify the increase in genetic gain due to the implementation of genomic selection within the traditional structure of a conventional breeding program. The design of these programs used the conventional program as a template. Minimal modifications were made to this template to produce the designs for each strategy.

#### Conventional Genomic Selection Breeding Program

The conventional plus genomic selection (ConvGS) breeding program used genomic selection to advance candidate DH lines in the testcross 1 and testcross 2 stages and to select parental lines for the subsequent breeding cycle. The parental lines were selected by choosing the 80 DH lines with the highest genomic estimated breeding values (GEBVs) from a set of candidates that comprised all DH lines from the testcross 2 stage and later yield trials. The minimum cycle time from bi-parental cross to the selection of new parental lines was 3 years, the same as in the conventional breeding program.

#### Genomic Selection Testcross 1 Breeding Program

The genomic selection testcross 1 (GS-TC1) breeding program used genomic selection to advance candidate DH lines in the testcross 1 and testcross 2 stages and to select parental lines for the subsequent breeding cycle. The parental lines were selected by choosing the 80 DH lines with the highest GEBVs from a set of candidates that comprised all DH lines from the testcross 1 stage and later yield trials. This reduced the minimum cycle time from bi-parental cross to the selection of new parental lines from 3 years in the conventional program to 2 years.

#### Genomic Selection Doubled Haploids Breeding Program

The genomic selection doubled haploids (GS-DH) breeding program used genomic selection to advance candidate DH lines in the testcross 1 and testcross 2 stages and to select parental lines for the subsequent breeding cycle. The parental lines were selected by choosing the 80 DH lines with the highest GEBVs from a set of candidates that comprised all DH lines from the DH stage. This reduced the minimum cycle time from bi-parental cross to the selection of new parental lines from 3 years in the conventional program to 1 year.

### Two-Part Breeding Programs

Crossing and selection of new parents in the two-part breeding programs was handled in the population improvement component (Fig. 3), which consisted of one or three crossing cycles per year (TP GS (1 cycle/year) and TP GS (3 cycles/year)). Parents were then grown in greenhouses and at the appropriate stage, crossings were undertaken following a circular scheme or a maximum avoidance of inbreeding scheme as described in the next section.

**Fig. 2.**
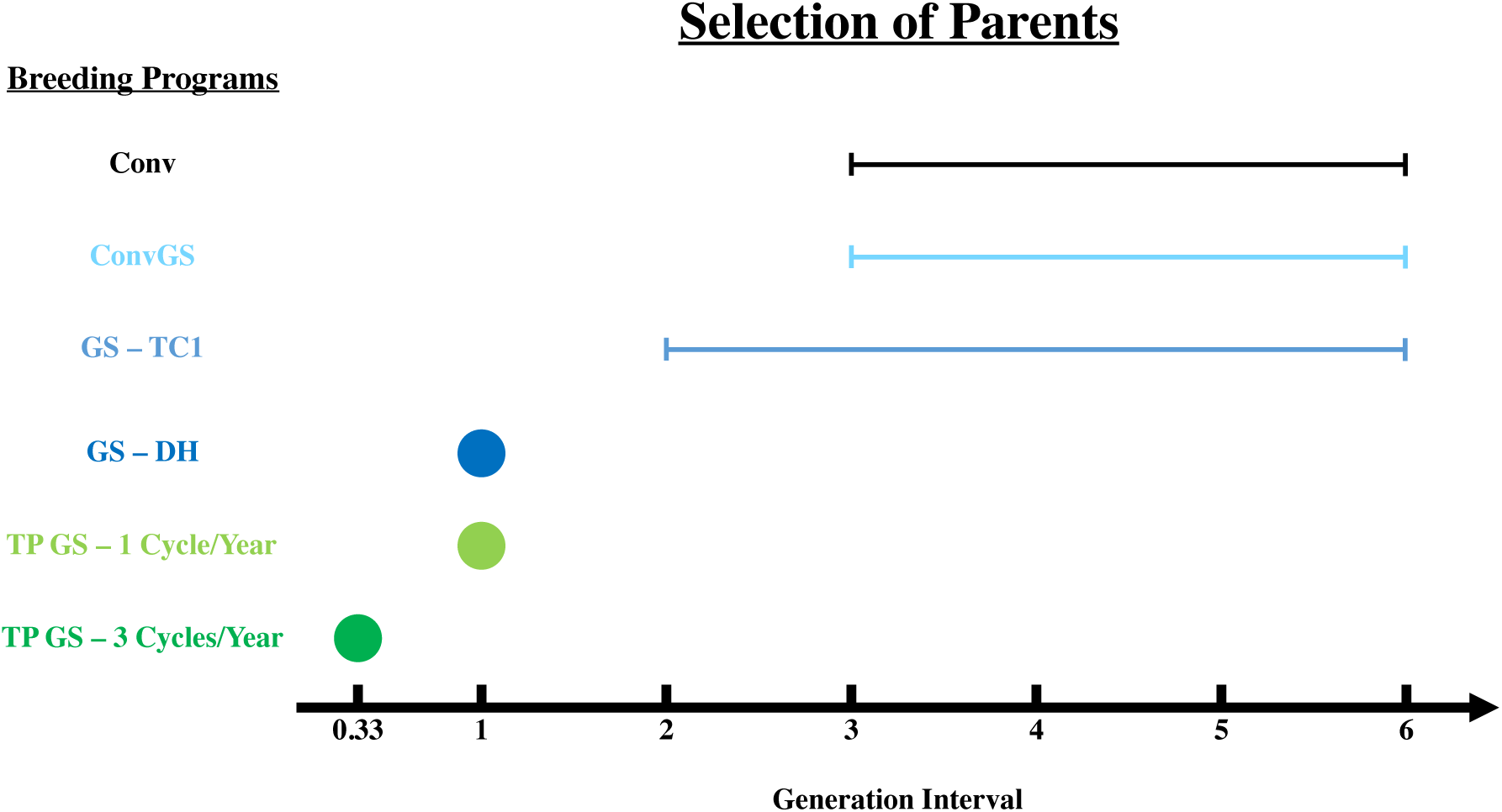
The generation intervals of hybrid crop breeding programs. Conv, conventional breeding program; ConvGS, conventional program with genomic selection; GS-TC1, genomic selection program with parents selected in the testcross 1 stage; GS-DH, genomic selection program with parents selected in the doubled haploid stage; TP GS, 1 Cycle/Year, two-part program with genomic selection; TP GS, 3 Cycles/Year, two-part program with genomic selection

**Fig. 3.**
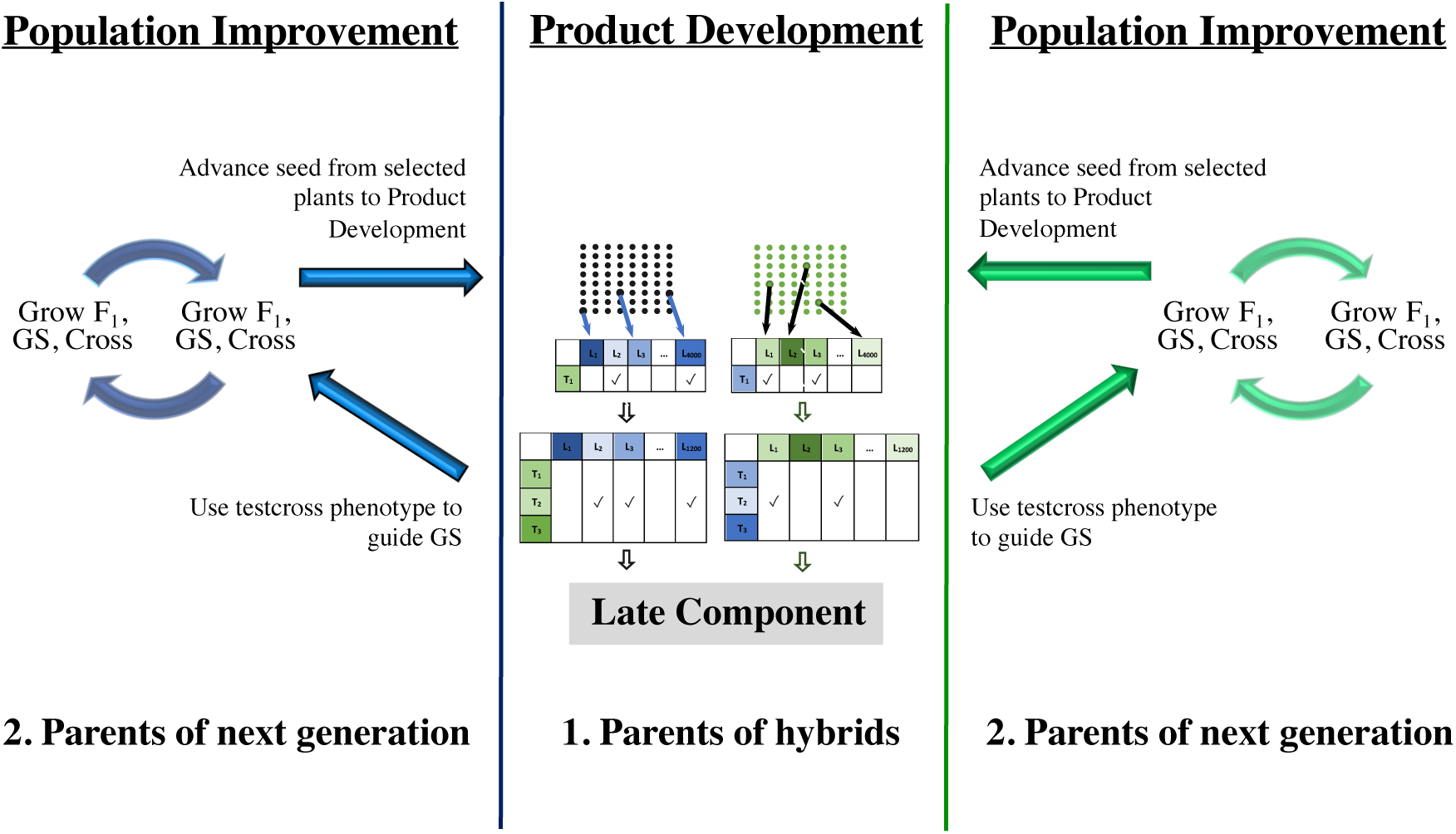
Overview of the two-part strategies with testcross genomic selection for hybrid crop breeding programs. (TP GS, 1 Cycle/Year, TP GS, 3 Cycles/Year). The number of crosses differed for each two-part breeding program to maintain equal operating costs. See Table 1

The product development component of the two-part program screened the germplasm to identify new commercial hybrids (Fig. 3). This process began with the production of the new DH lines. The DH lines were screened for testcross and single cross hybrid performance in the same manner as in the conventional plus genomic selection strategies. In two-part breeding programs none of the DH lines were selected for the crossing block, but their genomic and testcross phenotypic data were added to the genomic selection training population. This allowed the genomic selection model used in the population improvement component to be updated over time as new material was evaluated in the field.

### Crossing of Parents

Two crossing schemes were used in hybrid crop breeding programs:

i. a circular crossing scheme; and
ii. a maximum avoidance of inbreeding crossing scheme.

The circular crossing scheme used both between-family and within-family selection to select the 80 selection candidates with the highest genomic estimated breeding values (GEBVs). In each generation 80 crosses were made, ensuring that each parent was used only once as a male and a female. Consequently, this circular crossing scheme is different from the ‘circular design’ described by Kimura & Crow (1963), which only conducts within-family selection.

Hybrid crop breeding programs using genomic selection, with a generation interval of 1 year or less, also used the maximum avoidance of inbreeding crossing scheme. Maximum avoidance of inbreeding is a crossing scheme that maintains uniform contributions and inbreeding coefficients across all crosses (Wright 1921; Kimura and Crow 1963). The maximum avoidance of inbreeding crossing scheme mates the least related crosses in the first generation and maintains this crossing structure over generations. In the present study, the maximum avoidance of inbreeding scheme used within-family genomic selection to select the 2 selection candidates with the highest GEBVs per cross as new parents, giving 160 parents in total. In each generation 80 crosses were made, ensuring that each parent was used only once as a male or a female.

### Comparison of Breeding Programs

The performance of each breeding program was measured by comparing genetic gain and genetic variance of hybrids from the EYT stage. These hybrids were the crosses between all DH lines at the EYT stage from the two heterotic pools. The EYT stage was examined because it is the earliest stage in which all breeding programs evaluate DH lines for single cross hybrid performance. Genetic gain and genetic variance in the breeding programs were assessed by plotting mean and variance of true genetic values for hybrids at the EYT stage over time.

Accuracy of genomic prediction, defined in the next section, was assessed at the DH stage. The DH stage was examined because it allowed an assessment of the ability to rank all possible DH lines as parents of hybrids in the two heterotic pools. Accuracy of genomic predictions was also assessed in the population improvement components to estimate the accuracy of parent selection in the two-part breeding programs.

To aid in visualization, the mean values were centred at the mean value for lines in Year 0 for each replicate. Year 0 was defined as the last year of the burn-in phase. Direct comparisons between breeding programs for genetic gain, genetic variance and accuracy of genomic predictions were reported as ratios with 95% confidence intervals (95% CI). These ratios and 95% CI were calculated by performing paired Welch’s t-tests on log-transformed values from the 10 simulation replicates. The log-transformed differences and 95% CI from the t-test were then back-transformed to obtain ratios (Ramsey and Schafer 2002). All calculations were performed using R (R Development Core Team 2014).

#### Measurement of genomic selection accuracy

Accuracy of genomic prediction was defined as the correlation between the general combining ability (GCA) of DH lines and their GEBV. The GCA of an individual was calculated as the sum of all the average effects at the QTL weighted by the individuals genotype at these QTL. The average effects of alleles were calculated using allele frequencies from the corresponding population of DH lines in the other heterotic pool and the true simulated additive and dominance effects for each QTL. Therefore, the GCA of a DH line reflected its average performance as a parent in single cross hybrids when crossed to all DH lines from the other heterotic group.

## Results

Genomic selection increased the rate of genetic gain compared to phenotypic selection in hybrid crop breeding programs, mainly by reducing the generation interval. The two-part hybrid crop breeding program, with a generation interval of 0.33 years, produced the most genetic gain regardless of genotype-by-year variance. Genomic selection increased the selection accuracy compared to phenotypic selection in the early stages of hybrid crop breeding programs. There was a perfect rank correlation between the reduction in the generation interval and the reduction of genetic variance in hybrid crop breeding programs. Genomic selection reduced the efficiency of conversion of genetic variance into genetic gain compared to phenotypic selection. However, the use of the maximum avoidance of inbreeding crossing scheme slowed the reduction of genetic variance and increased the efficiency of conversion of genetic variance to genetic gain compared to the circular crossing scheme.

### Genetic Gain

Genomic selection increased the rate of genetic gain compared to phenotypic selection in hybrid crop breeding programs, mainly by reducing the generation interval. This is shown in Fig. 4, which presents the mean genetic value of hybrids at the elite yield trial stage. The first graph shows the trends for the mean for each of the breeding programs evaluated in the future breeding component when genotype-by-year variance is 0. The second graph shows the same trends for genotype-by-year variance of 4. Both graphs show that the two-part breeding program, with the shortest generation interval of 0.33 years, produced the most genetic gain. When genotype-by-year was 0, the TP GS (3 cycles/year) breeding program, which had the shortest generation interval of 0.33 years, generated 2.01 times the genetic gain of the Conv breeding program, which had the longest generation interval of 3 years.

**Fig. 4.**
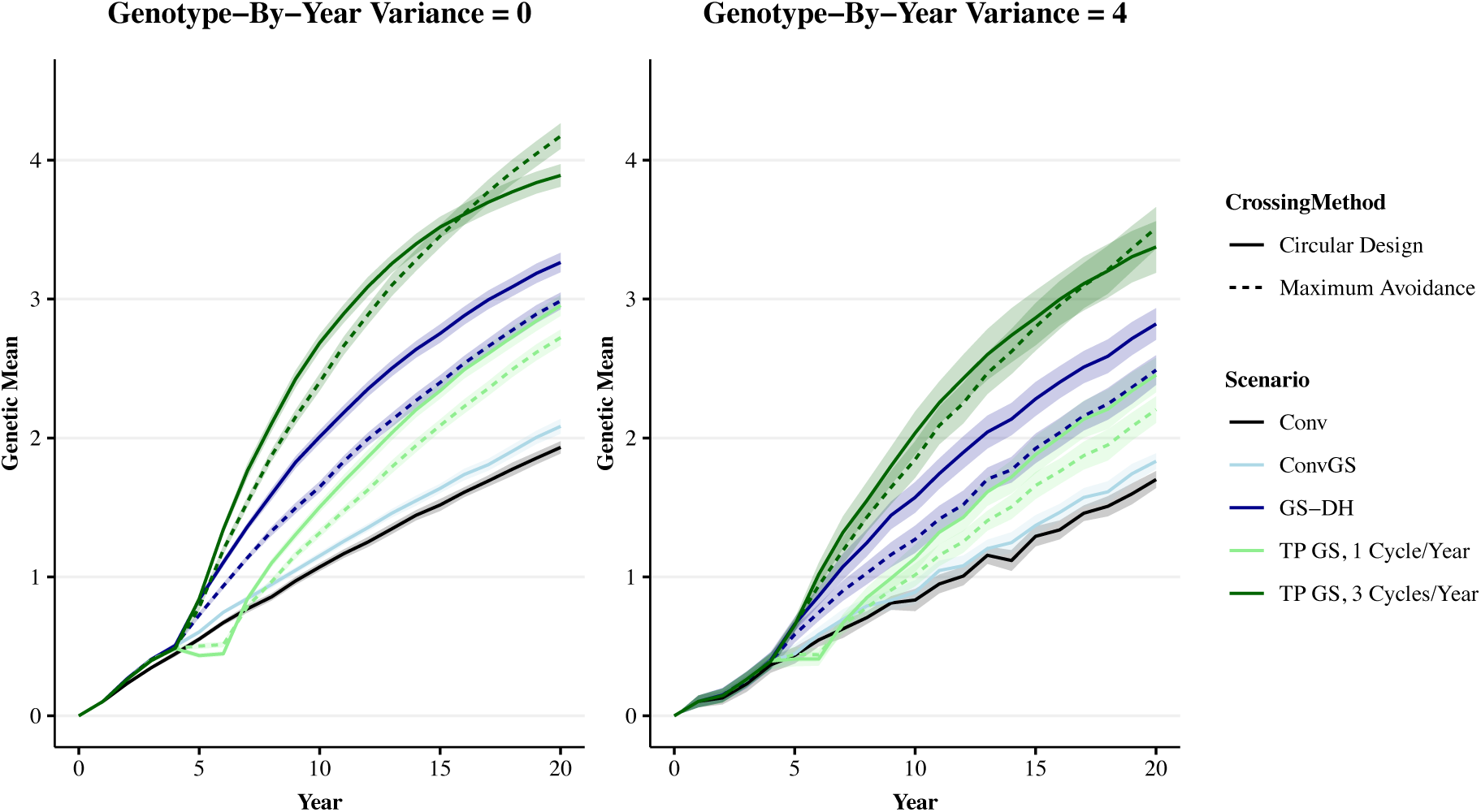
Genetic gain for all breeding programs over simulation years. Genetic gain when genotype-by-year variance was 0 and 4. Genetic gain is expressed as mean genetic value of hybrids at the elite yield trial stage over time. The mean genetic value for all replicates were centered on 0 in Year 4. Means for all 10 replicates are shown with dark lines, with the shaded area representing the 95% confidence intervals of the mean. Conv, conventional breeding program; Conv GS, conventional program with genomic selection; GS-DH, genomic selection program with parents selected in the doubled haploid stage; TP GS, 1 Cycle/Year, two-part program with genomic selection; TP GS, 3 Cycles/Year, two-part program with genomic selection

Fig. 4 also shows that the ranking of hybrid crop breeding programs for genetic gain was consistent across different genotype-by-year variances. This is shown by the average genetic values of hybrids in the final year (Year 20) of each graph. There was a perfect rank correlation between the generation interval and genetic gain of the breeding programs. Both graphs show the ranking from highest to lowest average genetic value was: TP GS (3 cycles/year), GS-DH, TP GS (1 cycle/year), ConvGS, Conv.

However, the relative differences between hybrid crop breeding programs using genomic selection and those using phenotypic selection were smaller when genotype-by-year variance was 4. At this level of genotype-by-year variance, the best performing two-part hybrid crop breeding program, TP GS (3 cycles/year), generated 1.96 times the genetic gain of the conventional breeding program. When genotype-by-year variance was 0, this value was 2.01. The relative differences in genetic gain between hybrid crop breeding programs using genomic selection remained constant across different levels of genotype-by-year variance.

Fig. 4 also shows that adding genomic selection to the conventional program without reducing the generation interval did not show a significant increase in genetic gain. This is shown by comparing genetic gain in the Conv program with genetic gain in the ConvGS program. The ConvGS program produced 1.08 (95% CI [1.00, 1.16]) and 1.08 (95% CI [0.98, 1.18]) times the genetic gain of the Conv program, when genotype-by-year variance was 0 and 4, respectively.

All breeding programs using genomic selection displayed a similar genetic gain prior to Year 5 (Fig. 4). Year 5 was the first year that hybrids at the elite yield trial stage were derived from parents selected by genomic selection. Therefore, the differences in genetic gain between Year 1 and Year 5 reflect the difference between using genomic selection or phenotypic selection on existing germplasm from the burn-in.

The TP GS (1 cycle/year) breeding program did not generate genetic gain in Year 5 and Year 6. This is because no selection was undertaken in the first two generations of future breeding of two-part hybrid crop breeding programs. These first two generations were required to convert the doubled haploid inbred parents from burn-in breeding into outbred parents. The TP GS (3 cycles/year) breeding program did not show this lag as it was able to complete this process and one cycle of selection within the first year of future breeding.

### Selection Accuracy

Genomic selection increased the selection accuracy compared to phenotypic selection in the early stages of hybrid crop breeding programs. This is show in Fig. 5, which plots the correlations between the simulated, true general combining abilities (GCA) for DH lines at the DH stage and their GEBV. The first graph shows the mean selection accuracy for all breeding programs when genotype-by-year variance was 0. The second graph shows the same trends for genotype-by-year variance of 4. The selection accuracies in the hybrid crop breeding programs using genomic selection were higher than those using phenotypic selection. In Year 1 when genotype-by-year variance was 0, all hybrid crop breeding programs using genomic selection had a selection accuracy of 0.73 while the Conv breeding program had a selection accuracy of 0.24.

**Fig. 5.**
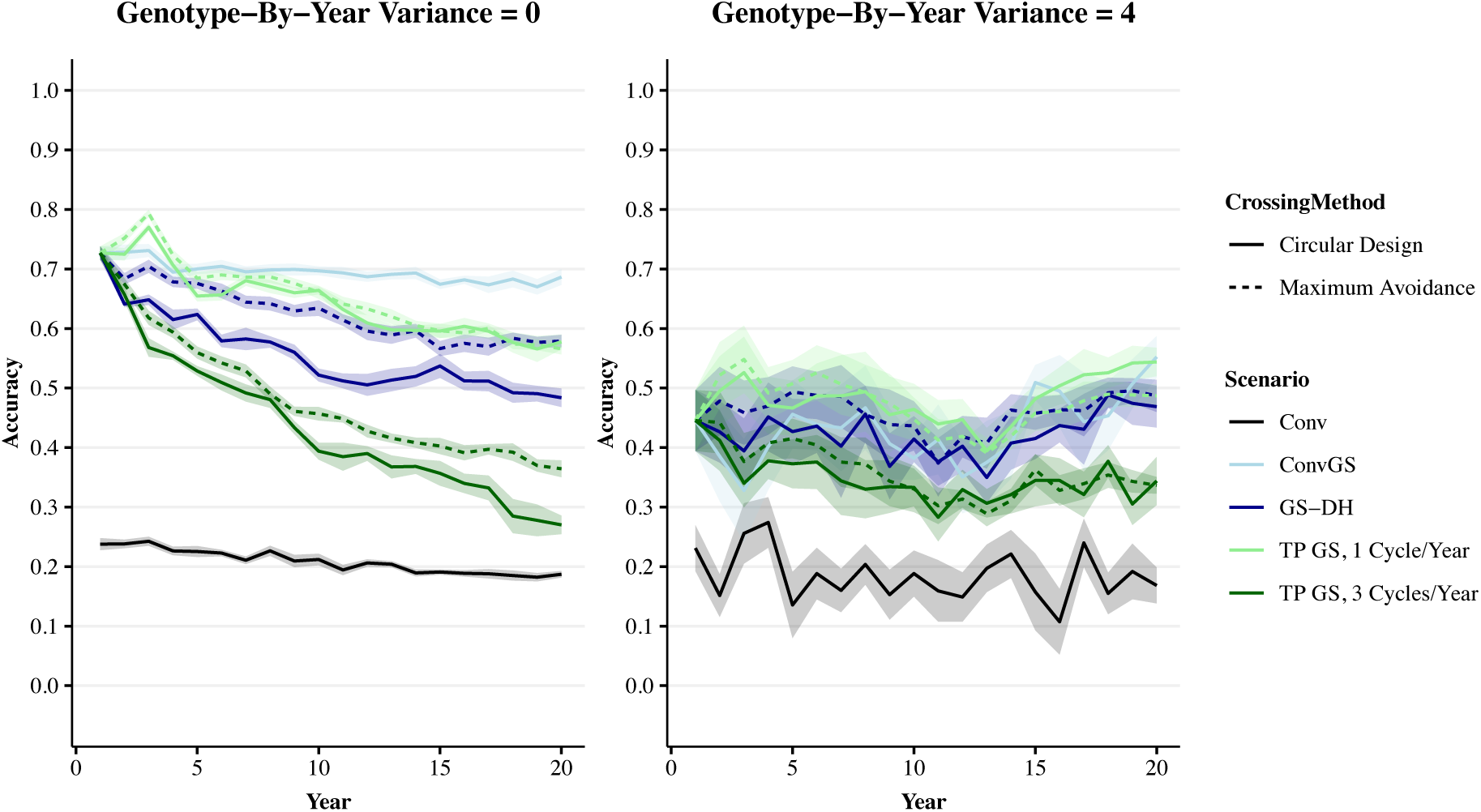
Selection accuracy for all breeding programs over selection cycles. Selection accuracy when genotype-by-year variance was 0 and 4. Selection accuracy is expressed as the correlation between true and predicted general combining abilities of doubled haploid (DH) lines at the DH stage over selection cycles. Means for all 10 replicates are shown with dark lines, with the shaded area representing the 95% confidence intervals of the mean. Conv, conventional breeding program; Conv GS, conventional program with genomic selection; GS-DH, genomic selection program with parents selected in the doubled haploid stage; TP GS, 1 Cycle/Year, two-part program with genomic selection; TP GS, 3 Cycles/Year, two-part program with genomic selection

Fig. 5 also shows that selection accuracies in all hybrid crop breeding programs decreased over the years of the simulation, and this decrease had a perfect rank correlation with the generation interval of hybrid crop breeding programs. This is shown by the average selection accuracy in the final year (Year 20) of each figure. Both figures show the ranking from highest to lowest average selection accuracy was: ConvGS, TP GS (1 cycle/year), GS-DH, TP GS (3 cycles/year). The ranking of hybrid crop breeding programs for selection accuracy was consistent across different genotype-by-year variances.

Selection accuracy in the two-part hybrid crop breeding programs was also measured in the population improvement stage. This is shown in Fig. 6, which plots the correlation between the simulated, true GCA for parental candidates and their GEBV for each cycle of crossing. There were one and three cycles of crossing per year. Therefore, each cycle is plotted at one third of a year increments in Fig. 6. The first figure shows the selection accuracy when genotype-by-year variance was 0. The second figure shows the selection accuracy when genotype-by-year variance was 4. Fig. 6 shows the change in selection accuracy over time differed between the two-part breeding programs. The first figure shows that the TP GS (1 cycle/year) breeding program displayed a gradual decrease in selection accuracy over time. The TP GS (3 cycles/year) breeding program displayed a faster decrease in selection accuracy and selection accuracy becomes 0 in Year 16. The second figure shows that selection accuracy remained constant over time when genotype-by-year was 4. Both graphs show yearly oscillations which correspond to the yearly updating of the training population.

**Fig. 6.**
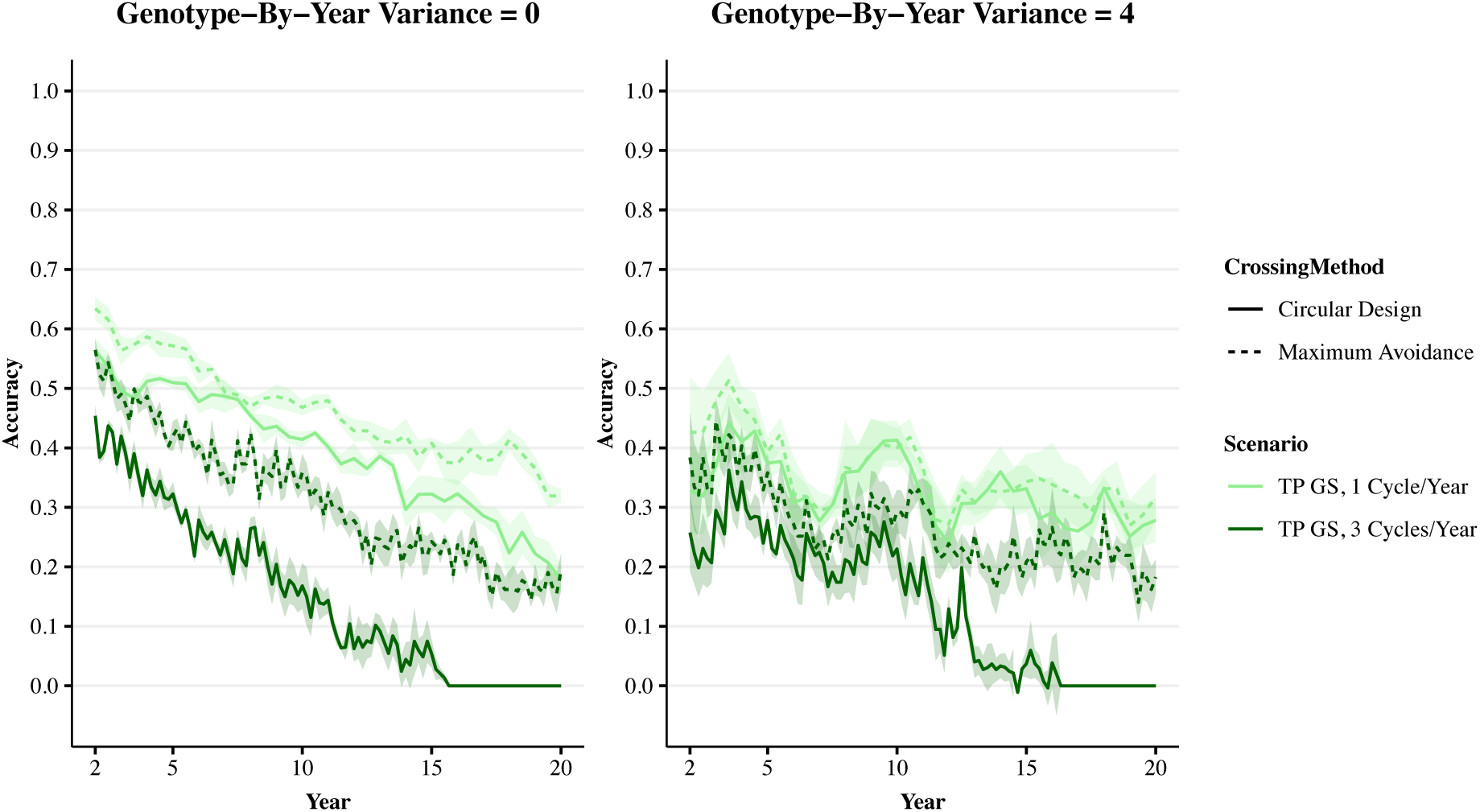
Within-family selection accuracy in the population improvement components of the two-part breeding programs. Within-family selection accuracy when genotype-by-year variance was 0 and 4. Within-family selection accuracy is expressed as the correlation between the simulated, true general combining abilities (GCA) for parental candidates and their predicted GCA. Means for all 10 replicates are shown with dark lines, with the shaded area representing the 95% confidence intervals of the mean. TP GS, 1 Cycle/Year, two-part program with genomic selection; TP GS, 3 Cycles/Year, two-part program with genomic selection

### Genetic Variance

All hybrid crop breeding programs displayed a reduction of genetic variance over simulation years. However, hybrid crop breeding programs using genomic selection caused a faster reduction of the genetic variance than the hybrid crop breeding program using phenotypic selection. This is shown in Fig. 7, which plots the change in genetic variance of hybrids at the elite yield trial stage over simulation years. The first graph shows the change in genetic variance for each future breeding program when genotype-by-year variance equals 0. The second graph shows the same breeding programs when genotype-by-year variance equals 4. The conventional breeding program displayed a gradual and consistent reduction of genetic variance across simulation years. All hybrid crop breeding programs using genomic selection displayed a large initial reduction of genetic variance. At Year 3 when genotype-by-year was 0, the Conv breeding program had 1.25 times the genetic variance of the ConvGS breeding program. The reduction of genetic variance in subsequent years differed between the hybrid crop breeding programs using genomic selection.

**Fig. 7.**
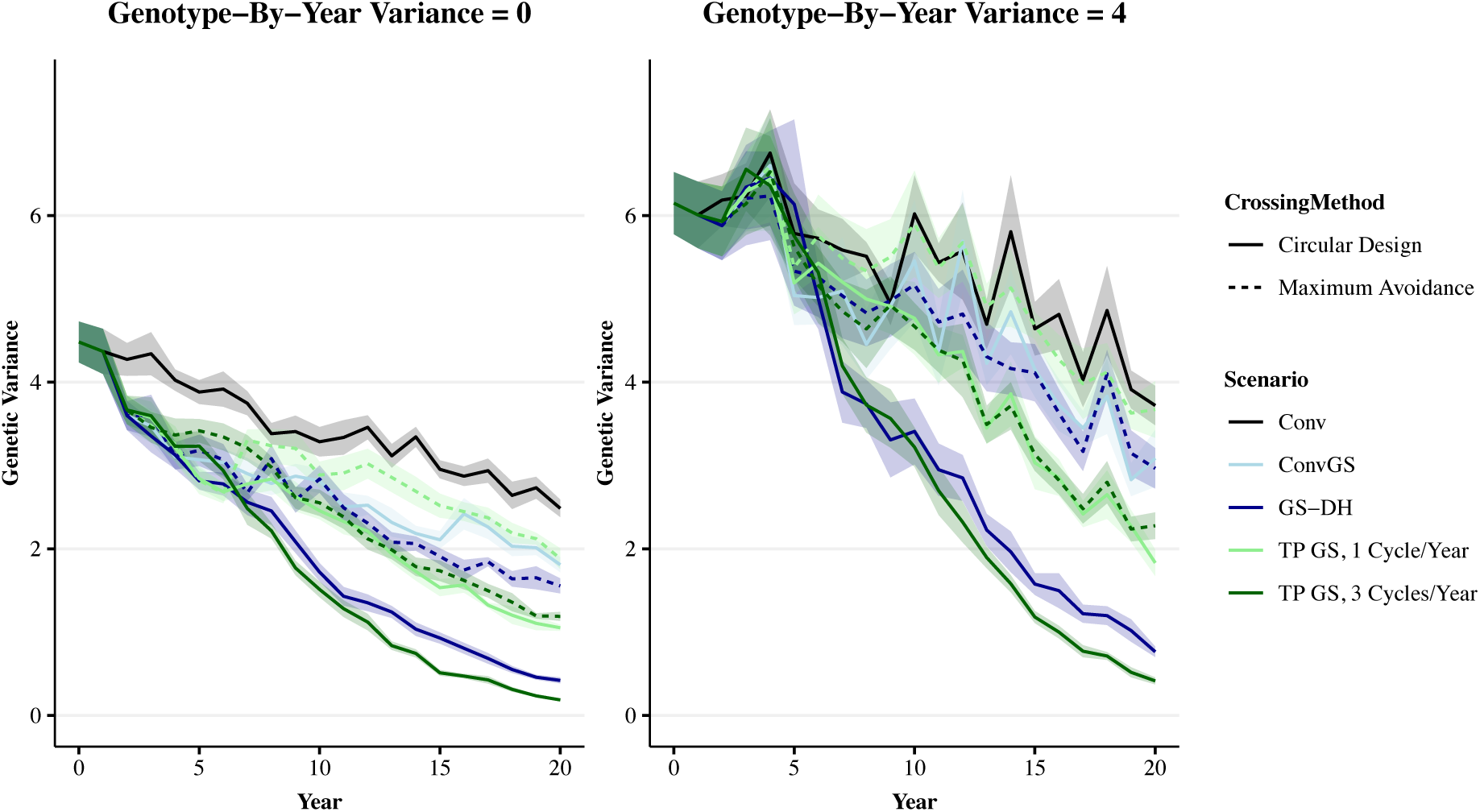
Genetic variance for all breeding programs over simulation years. Genetic variance when genotype-by-year variance was 0 and 4. Genetic variance is expressed as the genetic variance among hybrids at the elite yield trial (EYT) stage over simulation years. Means for all 10 replicates are shown with dark lines, with the shaded area representing the 95% confidence intervals of the mean. Conv, conventional breeding program; Conv GS, conventional program with genomic selection; GS-DH, genomic selection program with parents selected in the doubled haploid stage; TP GS, 1 Cycle/Year, two-part program with genomic selection; TP GS, 3 Cycles/Year, two-part program with genomic selection

Fig. 7 shows the change in genetic variance had a perfect rank correlation with the generation interval of hybrid crop breeding programs using genomic selection. This is shown by the genetic variance in the final year (Year 20) of each plot. Both graphs show the ranking from highest to lowest genetic variance was: ConvGS, TP GS (1 cycle/year), GS-DH, TP GS (3 cycles/year). When genotype-by-year variance was 0, the TP GS (3 cycles/year) breeding program had 0.44 times the genetic variance of the GS-DH breeding program and 0.07 times that of the Conv breeding program in Year 20. When genotype-by-year variance was 4, these values were 0.53 and 0.11, respectively. The ranking of hybrid crop breeding programs for genetic variance was consistent across different genotype-by-year variances.

### Crossing Schemes

Genomic selection with the maximum avoidance of inbreeding crossing scheme produced similar genetic gain over the circular crossing scheme, but maintained higher levels of genetic variance. Therefore, the maximum avoidance of inbreeding crossing scheme had a higher conversion efficiency compared to the circular scheme. This is shown in Fig. 8, which plots the genetic gain against the reduction of genetic variance for each hybrid crop breeding program-crossing scheme combination. All hybrid crop breeding programs using genomic selection with the circular crossing scheme had lower efficiency than the conventional breeding program using phenotypic selection. The first graph shows the trends for the mean values when genotype-by-year variance is 0. The second graph shows the same trends for genotype-by-year variance of 4. Fig. 8 also shows that hybrid crop breeding programs using genomic selection with the maximum avoidance crossing scheme had comparable conversion efficiency to the conventional breeding program using phenotypic selection. This ranking was consistent across different levels of genotype-by-year variance. Because the conversion efficiencies followed a very clear non-linear path (Fig. 8) it was not possible to formally test for statistically significant differences.

**Fig. 8.**
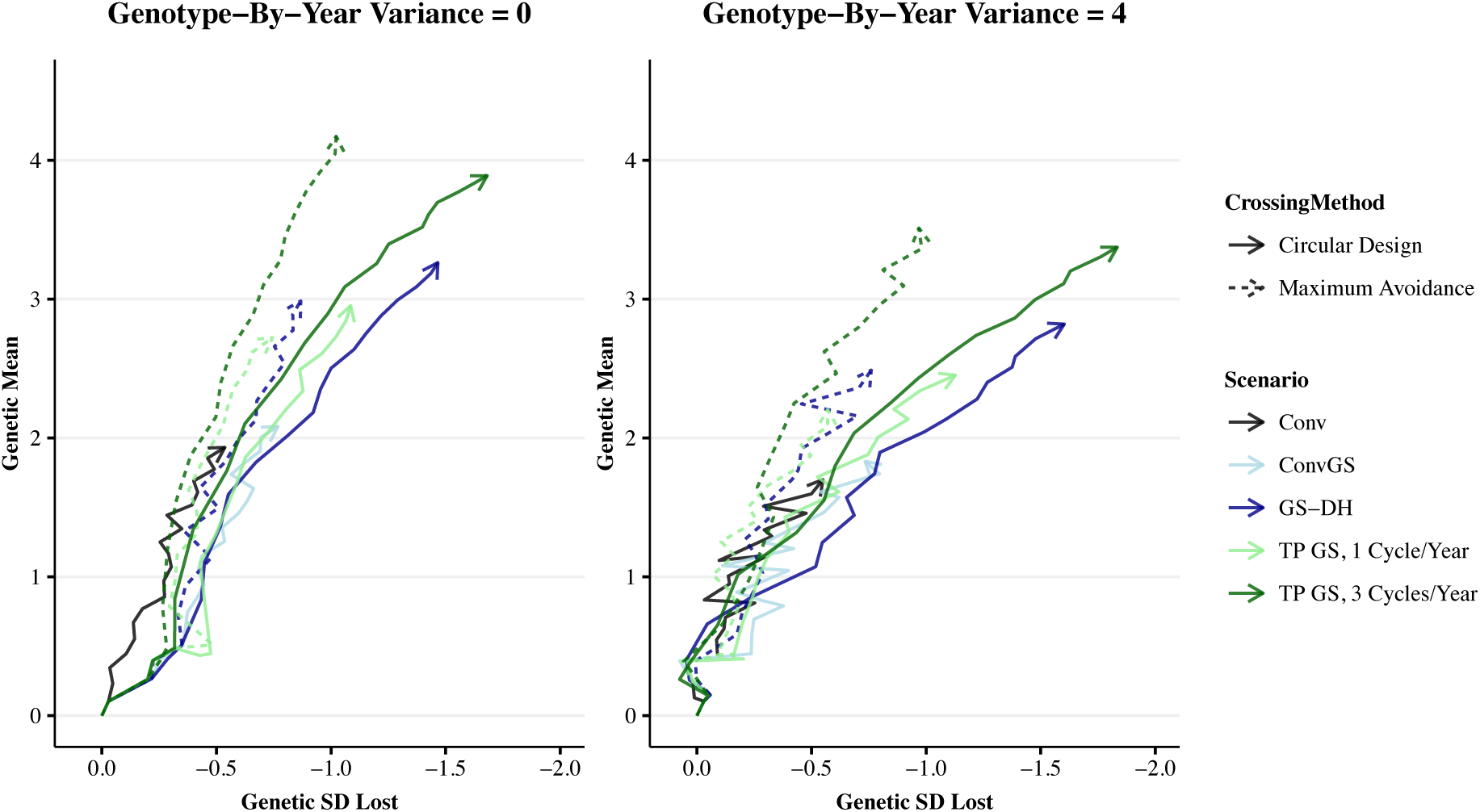
Conversion efficiency for all breeding programs over simulation years. Conversion efficiency when genotype-by-year variance was 0 and 4. Conversion efficiency is presented as the genetic gain against the genetic variance among hybrids at the elite yield trial between Year 0 and Year 20 of the simulation. Means for all 10 replicates are shown. Conv, conventional breeding program; Conv GS, conventional program with genomic selection; GS-DH, genomic selection program with parents selected in the doubled haploid stage; TP GS, 1 Cycle/Year, two-part program with genomic selection; TP GS, 3 Cycles/Year, two-part program with genomic selection

## Discussion

The results of this study highlight four points for discussion:

i. the impact of reciprocal recurrent genomic selection on the drivers of genetic gain in hybrid crop breeding programs;
ii. the impact of crossing schemes on genetic gain in hybrid crop breeding programs;
iii. the limitations of the simulation undertaken; and
iv. the practical implementation of the two-part strategy in real hybrid crop breeding programs.

### The impact of reciprocal recurrent genomic selection on the drivers of genetic gain in hybrid crop breeding programs

#### Generation Interval

Reciprocal recurrent genomic selection increased the rate of genetic gain in hybrid crop breeding programs, mainly by a reduction in generation interval. In an animal breeding context reciprocal recurrent genomic selection is a method that uses crossbred data to predict parent specific breeding values (Kinghorn et al. 2010). In a hybrid crop breeding context, parent specific breeding values can be predicted using hybrid phenotypes and inbred parental genotypes, and previous studies have predicted that reciprocal recurrent genomic selection could improve rates of genetic gain (Longin et al. 2013; Rembe et al. 2019). The results in the present study support this, showing that using reciprocal recurrent genomic selection increased genetic gain within a conventional hybrid crop breeding program design. However, the development of inbred parents takes time, which increases the generation interval (Griffing 1975). The recently proposed two-part strategy (Gaynor et al., 2017) provides a framework to remove this time delay by using outbred parents. In the context of hybrid crop breeding programs, the two-part strategy enables reciprocal recurrent genomic selection and the completion of multiple generations per year. The two-part strategy for hybrid crop breeding programs outlined in this study used this principle to achieve a generation interval of 0.33 years. Over the first ten years of breeding this drove a 1.33-fold increase in genetic gain compared to the best performing conventional plus genomic selection breeding program. While this is an important increase in rate of genetic gain, it is lower than the 3-fold expectation based on the breeder’s equation (Lush 1943), which can be explained by decreases in the selection accuracy and genetic variance over time.

#### Selection Accuracy

Rapid decreases in genomic selection accuracy were observed in two-part hybrid crop breeding programs and can be explained by:

i. a larger number of generations separating individuals in the ‘training population’ and selection candidates; and
ii. the reduction of the genetic variance of the selection candidates due to genomic selection.

In the present study, the two-part breeding program, with the shortest generation interval of 0.33 years, displayed the fastest decrease in genomic selection accuracy over time. The decreasing trend in selection accuracy is consistent with previous simulations of two-part breeding strategy for inbred crops (Gaynor et al. 2017; Gorjanc et al. 2018), although, the trends in the present study were much larger. There are a number of possible reasons for this.

Fluctuations in the average effect (of an allele substitution) over time driven by different trait genetic architectures can partly explain the lower selection accuracy in the present study compared to previous studies. The present study simulated a trait including both additive and dominance variation. When dominance contributes to trait variation, the average effects of alleles can change due to changes in allele frequencies (Falconer and Mackay 1996). The genomic estimated average effects of alleles are confounded with the ‘training population’ allele frequency, so they may not provide good estimates for average effects of alleles in the selection candidate whose allele frequency can be different. The previous studies of Gaynor et al. (2017) and Gorjanc et al. (2018) simulated strictly additive traits, so the average effects of alleles remained constant.

The more rapid decrease in genomic selection accuracy in this study compared to previous implementations of two-part strategies may also be partially attributable to different strategies for updating the ‘training population’. Previous studies accumulated records in the ‘training population’ across all years of future breeding, while in this present study, the size of the ‘training population’ was kept constant over time by updating the training population using a 3-year sliding window approach. This resulted in a training population that trended to half the size of that used in Gaynor et al. (2017) and a quarter of the size used in Gorjanc et al. (2018). Thirdly, the higher number of generations per year used in population improvement caused a greater divergence in relatedness between the training population and selection candidates compared to Gaynor et al. (2017).

#### Genetic Variance

The two-part strategy caused a rapid reduction of genetic variance in hybrid crop breeding programs. The reduction of genetic variance when using the two-part strategy may largely depend on how the additional genotyping costs in the population improvement are offset. Under a fixed budget, the additional genotyping costs could be offset in two simple ways. In the first strategy, the number of genotyped individuals per cross could be held constant while the number of parents and crosses could be reduced for each additional cycle of population improvement per unit time. However, such a strategy would cause a reduction of the effective population size and this could expose the breeding program to accelerated reduction of genetic variance due to genetic drift (Charlesworth 2009). Therefore, to overcome this risk, this study implemented an alternative strategy that maintained the number of parents and crosses but reduced the number of genotyped individuals per cross. This alternative strategy aims to maintain the effective population size and mitigate the reduction of genetic variance due to genetic drift. The results from the present study show that this alternative strategy was not sufficient to mitigate the accelerated reduction of genetic variance in two-part hybrid crop breeding programs using the circular crossing scheme. The two-part breeding program using the circular crossing scheme, with a generation interval of 0.33 years, displayed the fastest decrease in genetic variance over time. The large decreases in genetic variance with such an aggressive approach limited long-term genetic gain. Therefore, the maximum avoidance of inbreeding crossing scheme was explored.

### The impact of crossing schemes on the conversion efficiency in hybrid crop breeding programs

Reciprocal recurrent genomic selection with the maximum avoidance crossing scheme had the highest conversion efficiency, comparable to that of phenotypic selection. Genomic selection with the circular design crossing scheme had the lowest conversion efficiency. The higher conversion efficiency of the maximum avoidance crossing scheme was driven by a slower reduction of genetic variance over time compared to the circular scheme.

Crossing schemes are designed to manage genetic variance in breeding populations by avoiding the mating of closely related individuals. The circular scheme used in the present study does this by equalising the contributions of each cross. However, it can create higher levels of inbreeding compared to other crossing schemes because multiple parents could be selected from the same family if sufficient differences in the estimates of family means exist. Over generations this could result in large decreases in genetic variance due to genetic drift. Maximum avoidance of inbreeding is a crossing scheme that maintains uniform contributions and inbreeding coefficients across all crosses (Wright 1921; Kimura and Crow 1963). The maximum avoidance of inbreeding crossing scheme mates the least related crosses in the first generation and uses within-family truncation selection to choose new parents. In the present study, maximum avoidance worked well. Two-part hybrid crop breeding programs using maximum avoidance showed much smaller reductions in genetic variance than those using circular scheme.

The higher conversion efficiency of the maximum avoidance crossing scheme was important for the long-term genetic gain of hybrid crop breeding programs. Long-term genetic gain is dependent on the ability to exploit the within-family component of a breeding value, which is called the Mendelian sampling term (Wray and Thompson 1990; Meuwissen 1997; Pong-Wong and Woolliams 1998; Woolliams et al. 2015). Reciprocal recurrent genomic selection enabled a high within-family selection accuracy (Fig. 8), which both the maximum avoidance and circular scheme crossing schemes exploit. However, the maximum avoidance scheme preserved more genetic variance which also maintained a higher within-family selection accuracy. Therefore, the two-part hybrid crop breeding programs using the maximum avoidance crossing scheme displayed the highest long-term genetic gain.

### The limitations of the study

The simulations conducted in the present study did not model the full complexity of actual hybrid crop breeding programs. In this section the limitations and impact of key assumptions are discussed:

i. assumptions about the genetic architecture;
ii. assumptions that impact genomic selection accuracy;
iii. assumptions about the reciprocal recurrent genomic selection model;
iv. assumptions about the making of crosses;
v. assumptions about the ratio between genotyping and phenotyping costs;
vi. assumptions about the complexity of the breeding goal;
vii. assumptions that impact the maintenance of genetic variance.

#### Assumptions about the genetic architecture

The simulated trait was controlled by 3,000 QTL, a dominance degree of 0.9, a dominance variance of 0.3, and a heterotic pool split that occurred 100 generations ago. These values were chosen as they produced long-term trends for inbred and hybrid performance that reflected those observed in real data (Troyer and Wellin 2009). The main focus of the present study was on genetic gain, which relates to general combining ability (Sprague and Tatum 1942). We hypothesise that tuning the parameters to match long term genetic gain results in greater uncertainty in variance due to specific combining ability rather than variance due to general combining ability. Therefore, the assumptions about the genetic architecture of the simulated trait are likely to have limited impact on the conclusions of this study.

#### Assumptions that impact genomic selection accuracy

The genomic selection accuracies observed in these simulations are likely higher than those in real-world conditions. As previously described by Gaynor et al. (2017), this was because of conditions in the simulation that favoured high genomic selection accuracy such as

i. molecular markers with no genotyping errors;
ii. genetic control of the trait that did not involve epistasis; and
iii. a closed breeding program.

The accuracy of genomic selection affects the genetic gain of the simulated breeding programs. These effects should affect all hybrid crop breeding programs using genomic selection similarly, which suggests that using the two-part strategy should still outperform the other genomic selection breeding strategies. However, the relative performance of breeding programs using phenotypic selection to the breeding programs using genomic selection could change. If this were to occur, the hybrid crop breeding programs using the two-part strategy should still outperform the conventional breeding program because of the magnitude of difference observed in the simulation.

#### Assumptions about the making of crosses

An important assumption was how the present study performed crossing. As in Gaynor et al. (2017), the present study did not consider maturity differences between male and female parents. Maturity differences between prospective parents could result in particular crosses being missed, which could bring additional costs. Maturity differences could have specific impacts on the two crossing schemes used in the present study. Maturity differences could prevent each parent from being used twice, once as a male and once as a female, which was assumed in the circular design crossing scheme. Maturity differences could also result in missed crosses in the maximum avoidance crossing scheme. However, the flexibility of the maximum avoidance crossing scheme could account for maturity differences between prospective parents by replacing them by their next best-ranking siblings. Such maturity differences could have a more substantial impact on two-part breeding programs compared to the conventional plus genomic selection breeding programs, due to a higher number of crosses per year and lower seed availability in the two-part. However, the implications of maturity differences are likely to be relatively small and therefore have a relatively small effect on the performance of hybrid crop breeding programs.

#### Assumptions about the ratio between genotyping and phenotyping costs

The present study considered the costs of genotyping and phenotyping to be equal, and may not reflect current or future cost ratios for different breeding operations. The ratio between genotyping and phenotyping costs is more than likely to reduce in the future. Improvements in technology and the benefits from economy of scale could reduce genotyping costs in the future. Phenotyping costs could also reduce in the future, but likely at a slower rate. Any reduction in the ratio between genotyping and phenotyping costs would reduce the reallocation of resources required for the deployment of genomic selection, which is likely to favour hybrid crop breeding programs with high genotyping requirements, such as those using the two-part strategy.

#### Assumptions about the complexity of the breeding goal

The commercial products of hybrid crop breeding programs are a small number of hybrid varieties that are each planted on thousands to millions of acres. Therefore, these commercial products need to meet a wide range of requirements across a wide range of potential target production environments. Hybrid crop breeding programs must consider multiple traits relating to agronomic performance, disease resistance, and end-use quality. The hybrid crop breeding programs examined in this simulation only considered a single quantitative trait with 3,000 QTL. We assumed that this trait represented grain yield. However, it could equally represent a selection index with a few additional assumptions: all traits are measured on all individuals, all traits are pleiotropic, and economic merit is linear.

#### Assumptions that impact the maintenance of genetic variance

The present study showed there might be a potential risk for the rapid reduction of genetic variance in two-part hybrid crop breeding programs. The present study may overemphasise the issue of reduction of genetic variance in hybrid crop breeding programs, due to a combination of simulating a closed system, non-epistatic trait architecture, high genomic selection accuracy and a simplified breeding goal. Two crossing schemes to mitigate the reduction of genetic variance, the circular design and maximum avoidance, were used in the present study. Still, the two-part hybrid crop breeding programs displayed rapid reductions of genetic variance using both crossing schemes. Optimal contribution selection (Meuwissen 1997; Kinghorn et al. 1999; Woolliams et al. 2015; Gorjanc et al. 2018) and optimal cross selection (Allier et al. 2019) enable more complex crossing designs to avert the reduction of genetic variance in breeding programs. Such methods have the added benefit of balancing the choice between the maintenance of genetic variance, versus genetic gain and can, therefore, be tailored to prioritise short- and long-term genetic gain in two-part hybrid crop breeding programs. However, due to the large number of simulated scenarios optimal contribution methods were not used because they have high computational costs compared to the crossing schemes used in the present study.

### Further opportunities for the two-part strategy in real hybrid crop breeding programs

The present study demonstrated that the deployment of the two-part strategy could significantly increase genetic gain compared to current hybrid crop breeding program designs. However, the implementation of the two-part strategy in hybrid crop breeding programs requires some additional developments.

Alternative strategies for managing genetic variation need to be explored. The maximum avoidance of inbreeding crossing scheme used in this simulation only works in a closed breeding pipeline, so it cannot accommodate germplasm exchange. The maximum avoidance of inbreeding crossing scheme also offers little freedom to alter the balance between genetic gain and maintenance of diversity. More advanced strategies based on optimal contribution selection offer the potential to address both these limitations and should be explored (Meuwissen 1997; Woolliams et al. 2015). Further, a pre-breeding process could be integrated into the population improvement component of each heterotic pool to introgress external germplasm from gene banks or other sources (Gorjanc et al. 2016; Yang et al. 2019).

The creation of an outbred training population could partially mitigate the rapid reduction of genomic selection accuracy in two-part hybrid crop breeding programs. The entries in this outbred training population would consist of progeny from testcrosses to genotyped plants from the population improvement component. The progeny would be evaluated in plots to test the merit of their parents. This strategy is similar to older strategies for early testing of inbred lines (Sprague 1946). This data could be used to increase selection accuracy because it would reduce the genetic distance between the training and prediction individuals, which is known to be a significant determinant of accuracy (Habier et al. 2007; Clark et al. 2012). For example, in the current simulation the maximum number of generations between the selection candidates and the most recent training population lines could be cut in half by bypassing the creation of DH lines. Finally, a more speculative additional use of such an outbred training population would be to directly derive inbred lines from this population via apomixis.

The further opportunities for the two-part strategy in hybrid crop breeding programs, outlined here, incur further costs such as additional genotyping. Therefore, the implementation of the two-part strategy requires resource reallocation to account for these additional costs. The present study reallocated resources by reducing the number of selection candidates evaluated in the product development pipeline and maximised the number of crosses per generation in the population improvement component. However, the optimal resource reallocation strategy may differ between breeding programs. For example, breeding programs serving a smaller geographical region could reduce the number of trial locations in the product development pipeline. Therefore, optimal resource reallocation strategies require further research.

The deployment of the two-part strategy is a fundamental change to current practices in breeding programs and is currently untested empirically. Therefore, the development of bespoke transition strategies is required which would build the data sets the two-part strategy requires and build empirical confidence in its performance.

## Conclusions

Hybrid crop breeding programs using a two-part strategy produced the most genetic gain, but a maximum avoidance of inbreeding crossing scheme was required for it to increase long-term genetic gain. The two-part strategy uses outbred parents to complete multiple generations per year in hybrid crop breeding programs. In contrast conventional plus genomic selection strategies are limited in this regard by the time they take to develop inbred lines. The maximum avoidance of inbreeding crossing scheme manages genetic variance by maintaining uniform contributions and inbreeding coefficients across all crosses. This study performed stochastic simulations to quantify the potential of a two-part strategy in combination with two crossing schemes to increase the rate of genetic gain in hybrid crop breeding programs. Three main conclusions can be drawn from the results:

i. the implementation of genomic selection in hybrid crop breeding programs increases the rate of genetic gain;
ii. the two-part strategy was the most cost-effective strategy for implementing genomic selection in hybrid crop breeding programs.
iii. two-part hybrid crop breeding programs completing multiple selection cycles per year should use crossing schemes to manage genetic variance.

As well as the benefits outlined in this study, the flexibility of the two-part strategy offers further opportunities to integrate new technologies to further increase genetic gain in hybrid crop breeding programs, such as the use of outbred training populations. However, the practical implementation of the two-part strategy will require the development of bespoke transition strategies to fundamentally change the data, logistics, and infrastructure that underpin hybrid crop breeding programs.

## Author contributions statement

OP and JMH conceived the study. OP, JMH and RCG designed the study. OP developed the plant breeding program simulation. OP wrote the manuscript with input from all authors. All authors read and approved the final manuscript.

## Acknowledgments

The authors acknowledge the financial support from the BBSRC ISPG to The Roslin Institute (BBS/E/D/30002275). This work has made use of the resources provided by the Edinburgh Compute and Data Facility (ECDF) (http://www.ecdf.ed.ac.uk).

## Conflict of interest

The authors declare that they have no conflict of interest.

## Notes

### Competing Interest Statement

The authors have declared no competing interest.

## References

Alexandratos N, Bruinsma J (2012) World agriculture towards 2030/2050. Land Use Policy 20:375. https://doi.org/10.1016/S0264-8377(03)00047-4

Allier A, Lehermeier C, Charcosset A, et al (2019) Improving Short- and Long-Term Genetic Gain by Accounting for Within-Family Variance in Optimal Cross-Selection. Front Genet 10:1006. https://doi.org/10.3389/fgene.2019.01006

Bernardo R (2014) Essentials of Plant Breeding. Stemma Press

Bernardo R, Yu J (2007) Prospects for Genomewide Selection for Quantitative Traits in Maize. Crop Science 47:1082. https://doi.org/10.2135/cropsci2006.11.0690

Charlesworth B (2009) Fundamental concepts in genetics: Effective population size and patterns of molecular evolution and variation. Nature Reviews Genetics 10:195–205. https://doi.org/10.1038/nrg2526

Chen GK, Marjoram P, Wall JD (2009) Fast and flexible simulation of DNA sequence data. Genome Res 19:136–142. https://doi.org/10.1101/gr.083634.108

Clark S a, Hickey JM, Daetwyler HD, van der Werf JH (2012) The importance of information on relatives for the prediction of genomic breeding values and the implications for the makeup of reference data sets in livestock breeding schemes. Genetics Selection Evolution 44:4. https://doi.org/10.1186/1297-9686-44-4

Duvick DN, Smith JSC, Cooper M (2010) Long-Term Selection in a Commercial Hybrid Maize Breeding Program. In: Janick J (ed) Plant Breeding Reviews. John Wiley & Sons, Inc., Oxford, UK, pp 109–151

Falconer DS, Mackay TFC (1996) Introduction to Quantitative Genetics. Longman, Harlow, UK

Gaynor RC, Gorjanc G, Bentley AR, et al (2017) A two-part strategy for using genomic selection to develop inbred lines. Crop Science 57:2372–2386. https://doi.org/10.2135trcropsci2016.09.0742

Gaynor RC, Gorjanc G, Wilson D, Hickey JM (2019) AlphaSimR. Version 0.11.0URL http://CRAN.R-project.org/package=AlphaSimR

Geiger HH, Gordillo GA (2009) DOUBLED HAPLOIDS IN HYBRID MAIZE BREEDING. Maydica 485–499

Gorjanc G, Gaynor RC, Hickey JM (2018) Optimal cross selection for long-term genetic gain in two-part programs with rapid recurrent genomic selection. Theor Appl Genet 131:1953–1966. https://doi.org/10.1007/s00122-018-3125-3

Gorjanc G, Jenko J, Hearne SJ, Hickey JM (2016) Initiating maize pre-breeding programs using genomic selection to harness polygenic variation from landrace populations. BMC Genomics 17:30. https://doi.org/10.1186/s12864-015-2345-z

Griffing B (1975) Efficiency changes due to use of doubled-haploids in recurrent selection methods. Theoret Appl Genetics 46:367–385. https://doi.org/10.1007/BF00281141

Habier D, Fernando RL, Dekkers JCM (2007) The impact of genetic relationship information on genome-assisted breeding values. Genetics 177:2389–2397. https://doi.org/10.1534/genetics.107.081190

Hull FH (1945) Recurrent selection and specific combining ability in corn. J Amer Soc Agron

Kimura M, Crow JF (1963) On the maximum avoidance of inbreeding. Genet Res 4:399–415. https://doi.org/10.1017/S0016672300003797

Kinghorn BP, Hickey JM, Werf JHJVD (2010) Reciprocal Recurrent Genomic Selection for Total Genetic Merit in Crossbred Individuals. Proc 9th WCGALP 0036

Kinghorn BP, Shepherd RK, Woolliams JL (1999). In: Proceedings of the Association for the Advancement of Animal Breeding and Genetics. p 412

Longin CFH, Gowda M, Mühleisen J, et al (2013) Hybrid wheat: quantitative genetic parameters and consequences for the design of breeding programs. Theor Appl Genet 126:2791–2801. https://doi.org/10.1007/s00122-013-2172-z

Lush JL (1943) Animal Breeding Plans. Iowa State Press

Meuwissen THE (1997) Maximizing the response of selection with a predefined rate of inbreeding. Journal of Animal Science 75:934–940. https://doi.org/doi:/1997.754934x

Pong-Wong R, A. Woolliams J (1998) Response to mass selection when an identified major gene is segregating. Genetics Selection Evolution 30:313–337

R Development Core Team (2014) R: A language and environment for statistical computing. R Foundation for Statistical Computing, Vienna, Austria

Ramsey FL, Schafer DW (2002) The Statistical Sleuth: A Course in Methods of Data Analysis, 2nd edn. Duxbury/Thomson Learning, Australia

Rembe M, Zhao Y, Jiang Y, Reif JC (2019) Reciprocal recurrent genomic selection: an attractive tool to leverage hybrid wheat breeding. Theor Appl Genet 132:687–698. https://doi.org/10.1007/s00122-018-3244-x

Sprague GF (1946) THE EXPERIMENTAL BASIS FOR HYBRID MAIZE. Biological Reviews 21:101–120. https://doi.org/10.1111/j.1469-185X.1946.tb00317.x

Sprague GF, Tatum LA (1942) General vs. Specific Combining Ability in Single Crosses of Corn 1. Agron J 34:923–932. https://doi.org/10.2134/agronj1942.00021962003400100008x

Troyer AF, Wellin EJ (2009) Heterosis decreasing in hybrids: Yield test inbreds. Crop Science 49:1969–1976. https://doi.org/10.2135/cropsci2009.04.0170

Whittaker JC, Thompson R, Denham MC (2000) Marker-assisted selection using ridge regression. Genet Res 75:249–252

Woolliams JA, Berg P, Dagnachew BS, Meuwissen THE (2015) Genetic contributions and their optimization. Journal of Animal Breeding and Genetics 132:89–99. https://doi.org/10.1111/jbg.12148

Wray NR, Thompson R (1990) Prediction of rates of inbreeding in selected populations. Genet Res 55:41–54. https://doi.org/10.1017/S0016672300025180

Wright S (1921) Systems of Mating. II. the Effects of Inbreeding on the Genetic Composition of a Population. Genetics 6:124–143

Yang CJ, Sharma R, Gorjanc G, et al (2019) Origin Specific Genomic Selection: a simple process to optimize the favourable contribution of parents to progeny. Genetics

